# Morphological phylogenetic analysis with inapplicable data

**DOI:** 10.1101/209775

**Authors:** Martin D. Brazeau, Thomas Guillerme, Martin R. Smith

**Author notes:** All authors contributed equally to this work.

## Abstract

Non-independence of characters is a real phenomenon in phylogenetic data matrices, even though phylogenetic reconstruction algorithms generally assume character independence. In morphological datasets, the problem results in characters that cannot be applied to certain terminal taxa, with this inapplicability treated as “missing data” in a popular method of character coding. However, this treatment is known to create spurious tree length estimates on certain topologies, potentially leading to erroneous results in phylogenetic searches. Here we present a single-character algorithm for ancestral states reconstruction in datasets that have been coded using reductive coding. The algorithm uses up to four traversals on a tree to resolve final ancestral states – which are required in full before a tree can be scored. The algorithm employs explicit criteria for the resolution of ambiguity in applicable/inapplicable dichotomies and the optimization of missing data. We score trees following a previously published procedure that minimizes homoplasy over all characters. Our analysis of published datasets shows that, compared to traditional methods, our new method identifies different trees as “optimal”; as such, correcting for inapplicable data may significantly alter the outcome of tree searches.

## Introduction

Morphological characters are an essential source of data in phylogenetic studies. Even though they have been outpaced in their use by molecular sequence data, they remain indispensable for a range of research programmes that depend on knowledge of extinct or ancestral phenotypic conditions (e.g. palaeontology, molecular clock calibrations, comparative developmental biology). Despite advances in the use of probabilistic models for analysing morphological data (e.g. Lewis, 2001; Wright et al., 2016), all transformation-based methods (e.g. parsimony, likelihood) are subject to the same persistent problem in morphological and phenotypic data: the logical inapplicability of characters.

Logical inapplicability occurs when a dataset contains characters that can only have a meaningful value in a subset the taxa under investigation. This usually arises when one or more characters are ontologically dependant on a neomorphic (presence/absence) character, here termed the “principal character”. The problems associated with coding these character relationships have been discussed in detail since the advent of desktop phylogenetic computer programs (Farris, 1988; Platnick et al., 1991; Maddison, 1993; Wilkinson, 1995; Pleijel, 1995; Strong and Lipscomb, 1999; Hawkins, 2000; Forey and Kitching, 2000; Fitzhugh, 2006; Brazeau, 2011) and reflect the mathematical consequences of several popular coding procedures (reviewed by Brazeau, 2011). As it stands, no existing software can accommodate the computational issues that arise from logical inapplicability.

For situations in which a the state of a character depends on the presence or absence of another character, there is widespread agreement that the best practice is to code inapplicable taxa using a token treated as “missing data”. The token “-” is often used to distinguish cases in which a character is inapplicable (e.g. tail colour, in a taxon that lacks a tail) from those in which a character’s state is uncertain (typically denoted “?” – e.g. tail colour, where the colour of the tail is unknown). For the tail example, this looks like:

- 1. Tail: absent (0); present (1)
- 2. Tail colour: blue (0); red (1); inapplicable (? or -)

This coding style, termed reductive or contingent coding (Strong and Lipscomb, 1999; Forey and Kitching, 2000, hereafter “reductive coding”), treats inapplicable state values as missing data – as though the characteristic in question is not preserved in known specimens. This approach is considered unlikely to lead to implicit (and unintended) character weighting, but does entail spurious calculations (Maddison, 1993): such a coding scheme will allow the reconstruction of transformations at nodes where the inferred state is logically impossible (e.g. a change in tail colour in an ancestor with no tail). These logically impossible state reconstructions and their concominant transformations have been informally referred to as “pseudo-parsimony”, but could be generalized to “pseudo-optimality”, since they would occur in probabilistic calculations as well. In spite of the problem of logically impossible state reconstructions, reductive coding is still widely used and defended (Strong and Lipscomb 1999; Hawkins 2000; Brazeau 2011; but see also arguments from Fitzhugh 2006 and Vogt 2017).

Maddison (1993) concluded that addressing this problem would require modification of phylogenetic software; 25 years later, there are still few signs of progress on this problem. Some recent and important theoretical advances were made by De Laet (2005, 2015), but De Laet does not describe a single-character algorithm; nor does he provide details of how his method might handle ambiguity, polymorphism, missing data, or multistate characters.

In this paper we detail modifications required to enable the Fitch algorithm to process morphological characters that exhibit inapplicability. We consider how trees should be evaluated for optimality, consistent with the method described by De Laet (2015). Furthermore, we show that the effect of “pseudo-optimal” reconstructions can lead to both significant over- and under-estimates of parsimony scores during tree searches. Our algorithm and its implementation allow a special token to indicate inapplicability, meaning that existing datasets that use the gap token to denote inapplicability can be treated with little modification. However, we show that investigators may wish to re-code some characters in ways that can avoid the inapplicable token altogether.

## Theoretical considerations of ancestral state reconstructions and tree scores

We wish to have an algorithm that: (i), incorporates all phylogenetically relevant information; (ii), generates logically and internally consistent nodal state sets, including the reconstruction of an inapplicable “state” where a character does not logically apply; and (iii), calculates exact optimality scores that neither over- nor under-penalize any given tree. In order to reconstruct ancestral states in an ontologically dependent character, it is first necessary optimize the presence or absence of the principal character, which dictates the nodes at which the ontologically dependent character is applicable. This opens up two questions: how do we resolve ambiguity in the principal character, and how do we calculate an optimality metric for the tree?

### Ancestral state reconstructions

It is not unusual for a character to have two mutually exclusive nodal reconstruction sets that are equally parsimonious. Particularly relevant here is the case of a presence/absence character whose distribution can be accommodated by one of two equally parsimonious explanations: the gain and loss of the character (accelerated transition / AccTran), or two parallel gains (delayed transition / DelTran).

By minimizing the number of independent origins in the neomorphic character, the AccTran optimization maximises the homology represented by that character. This is preferable – on the principle of maximising homology and minimising homoplasy (De Laet, 2005) – to the DelTran optimization (de Pinna, 1991), in which each independent gain of the character represents an additional instance of homoplasy.

Even though the neomorphic character makes an equal contribution to tree length under either reconstruction, the contribution of any ontologically dependent characters may depend on the optimization chosen. By way of example, an AccTran optimization that reconstructs the gain and loss of a red tail implies that all tails are homologous; there is therefore no homoplasy in the ontologically dependent character “tail colour”. In contrast, a DelTran optimization that invokes two parallel gains of a red tail on the same tree may be equally parsimonious with respect to the principal character, but would imply two independent origins of red colouration, one of which represents an instance of homoplasy.

A satisfactory method must therefore distinguish the presence of a character from its absence: something that is impossible within the Fitch algorithm, which is blind to whether tokens denote presence, absence, or some other property, and therefore cannot differentiate gain/loss from parallel gain. Thankfully, the presence or absence of a principal character is implicit in the distinction between applicable and inapplicable states in an ontologically dependent character. If a dataset is coded accurately, then the applicable/inapplicable distinction will exactly mirror the presence/absence distribution of its principal character. That means that knowledge of presence/absence can be built into the handling of ontologically dependent characters, and that the algorithm need not be explicitly supplied with a prior specification of the principal character.

### Scoring trees

For the purpose of phylogenetic searches, there must be some function for evaluating the amount of homoplasy implied by the optimal character reconstructions on a particular tree topology. When all characters are applicable to all terminal taxa, the amount of homoplasy is simply equal to the number of transformations minus the theoretical minimum number of transformations over all characters. However, a transformation between applicable and inapplicable states has no clear meaning with respect to length counts (i.e. it is not an *independent* transformational event).

The problem is most clearly illustrated in the context of a principal character with a number of ontologically dependent transformational characters (Fig. 1). If each transformation from an applicable to an inapplicable state contributes to tree length, the loss of the principal character will be severely penalised, even though it can be explained as a single evolutionary event. In contrast, if transformations between applicable and inapplicable states contribute nothing to tree length, then losses and gains of the principal character are inadequately penalised, effectively resulting in a penalty for character congruence, and thus a penalty for homology.

This illustrates why the amount of homoplasy within a tree cannot simply be expressed in terms of the number of transformational events when ontologically dependent characters are present (De Laet, 2005, 2015). To once again borrow Maddison’s (1993) example, a single transformation from “tail absent” to “tail present, red” does not represent an instance of homoplasy for the ontologically dependant character “tail colour”, but if this same transformation happens twice, homoplasy in tail colour has occurred: the tree should be penalized once for the independent origin of the second tail, and once more because the second tail, when it appeared, happened to exhibit the same state (red) as the first (Fig. 2). However, the loss of a tail implies the simultaneous loss of colour and other similar attributes, which cannot similarly be explained as transformations.

**Figure 1:**
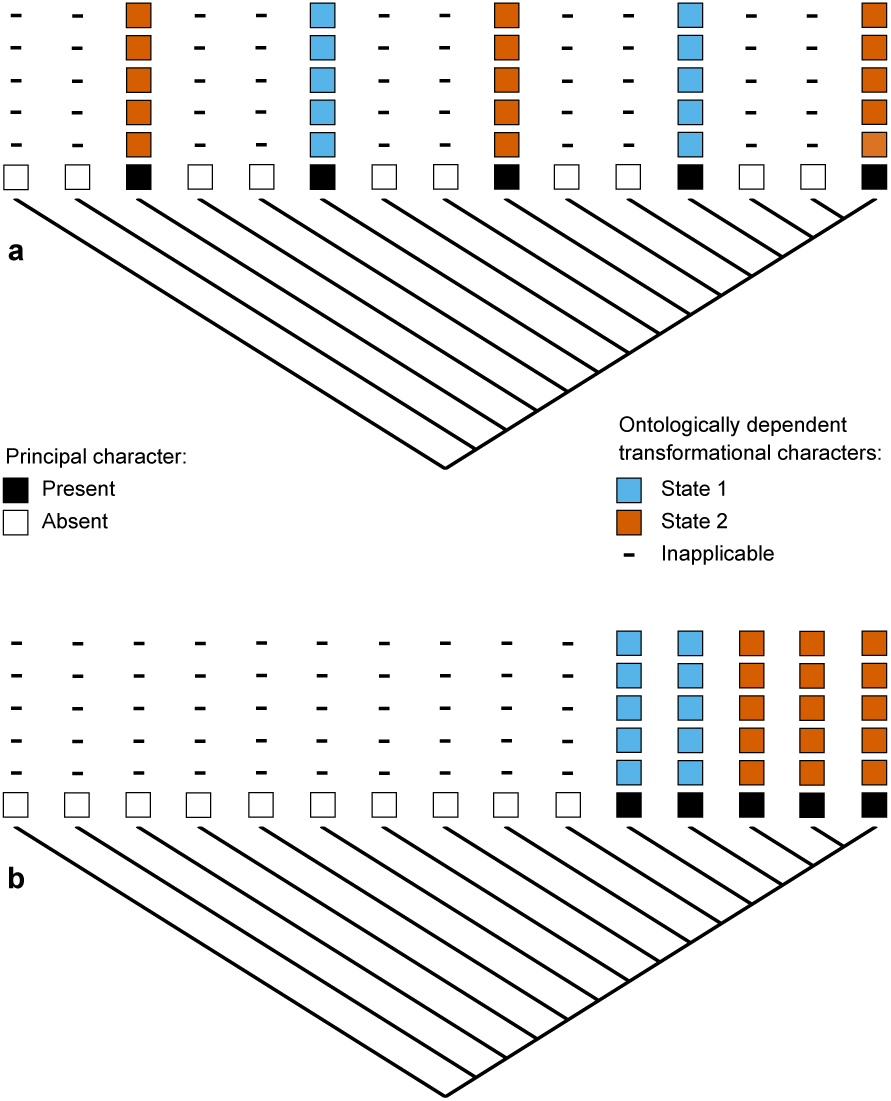
Effect of counting method on tree preference. If transformations between losses and gains of the principal character are inadequately penalised, then trees with multiple gains of the principal character (**a**) will be favoured; if transformations between applicable and inapplicable states are penalized, then trees in which the principal character evolves exactly once (**b**) will be favoured.

A satisfactory handling of inapplicable data in morphology must satisfy at least two criteria: (i), non-redundancy; and (ii), maximizing the explanatory value of the data (De Laet, 2005). This is not possible with currently implemented algorithms. De Laet (2005) proposes a solution in which the penalty on the tree is not simply the sum of steps, but also the number of regions (or “subcharacters”) defined by a character.

Regions are defined as subtrees in which a character is logically applicable (i.e. applicable character regions). De Laet proposes that the optimal tree is that which minimizes the sum of the number of regions and the number of transformations between states.

Throughout this manuscript we therefore make a clear distinction between tree length and tree score, either of which might be chosen as an optimality criterion. Tree length designates the number of transformational events (steps) implied by a topology, whereas tree score designates an optimisation value that combines some function of the tree length with other non-transformational events, such as the sum of the number of regions.

## Fitch parsimony with partially applicable characters

The algorithm described below is a single-character (*sensu* Ronquist, 1998) method in which “inapplicability” is reserved as a special token (usually denoted with the symbol “-”). At its core, the algorithm resembles the Fitch algorithm (Fitch, 1971), with “inapplicability” being treated as an additional state. The first two passes of the algorithm use the distribution of applicable and inapplicable tokens to infer whether the associated principal character can be optimally reconstructed as present at each node. In nodes where the principal character can be reconstructed as present, normal Fitch rules are used to identify and count transformations. Transformations are not reconstructed at nodes where the principal character was necessarily absent.

**Figure 2:**
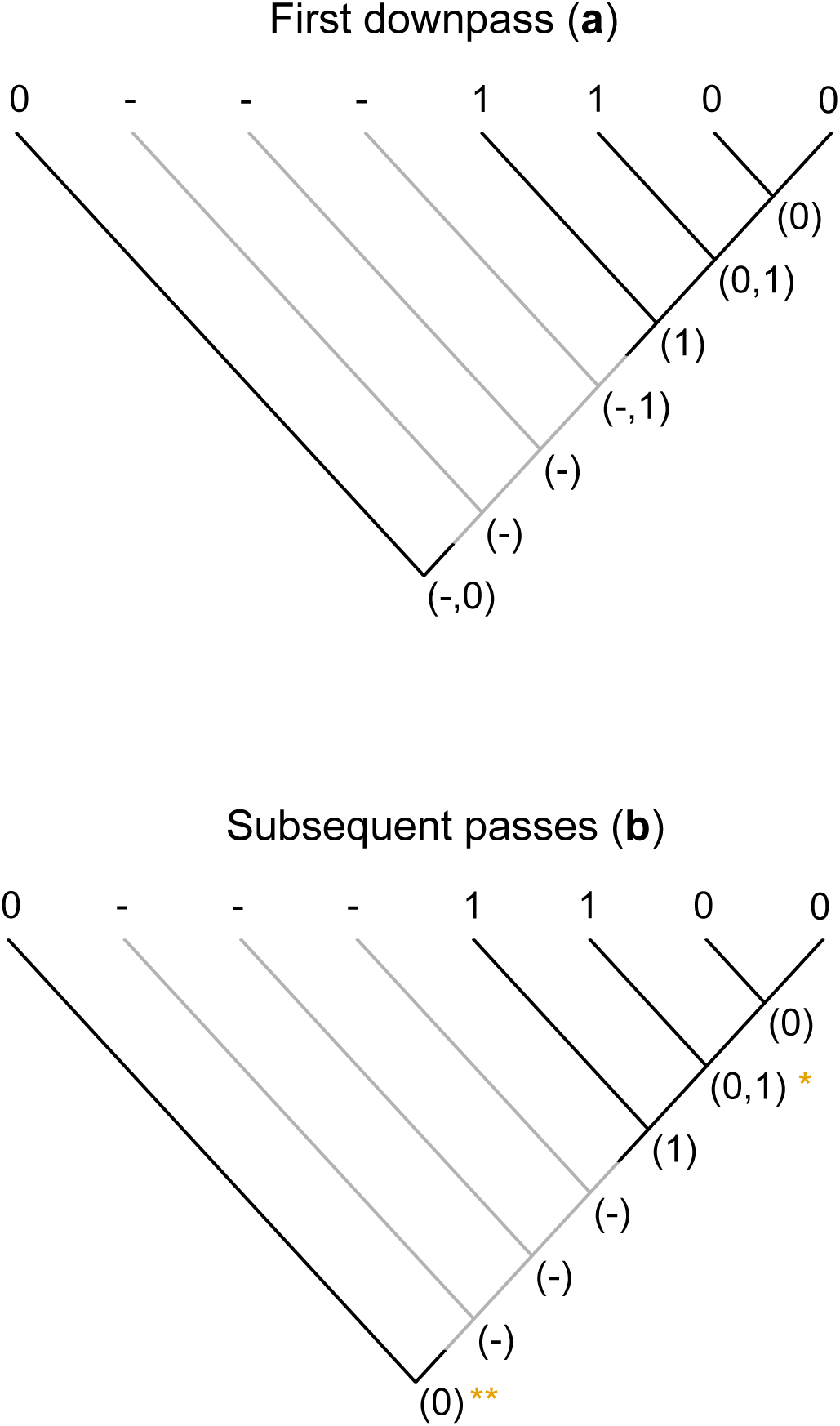
Scoring of a simple tree with inapplicable data. A principal character is present in two regions of the tree (black lines). A transformation from state 1 to state 0 adds one step to tree length. A second occurrence of state 0 represents a case of homoplasy, and should also contribute to tree score. In our algorithm, the first downpass (a) generates possible reconstructions of each node; final state reconstructions are generated in the first uppass and not modified by further passes. The second downpass increments tree length by two, reflecting one step (at *) ^11^ additional region (at **).

To count the number of applicable regions, a flag is stored at each node which records whether or not any descendants store applicable values. This allows the number of regions to be incremented when moving, on the second downpass or on the second uppass, from an inapplicable region of the tree to an applicable region. Up to four passes are therefore required to complete ancestral state reconstructions for a given node (Fig. 3). An interactive visualisation of the four passes is available via the Inapp R package at https://github.com/TGuillerme/Inapp.

**Figure 3:**
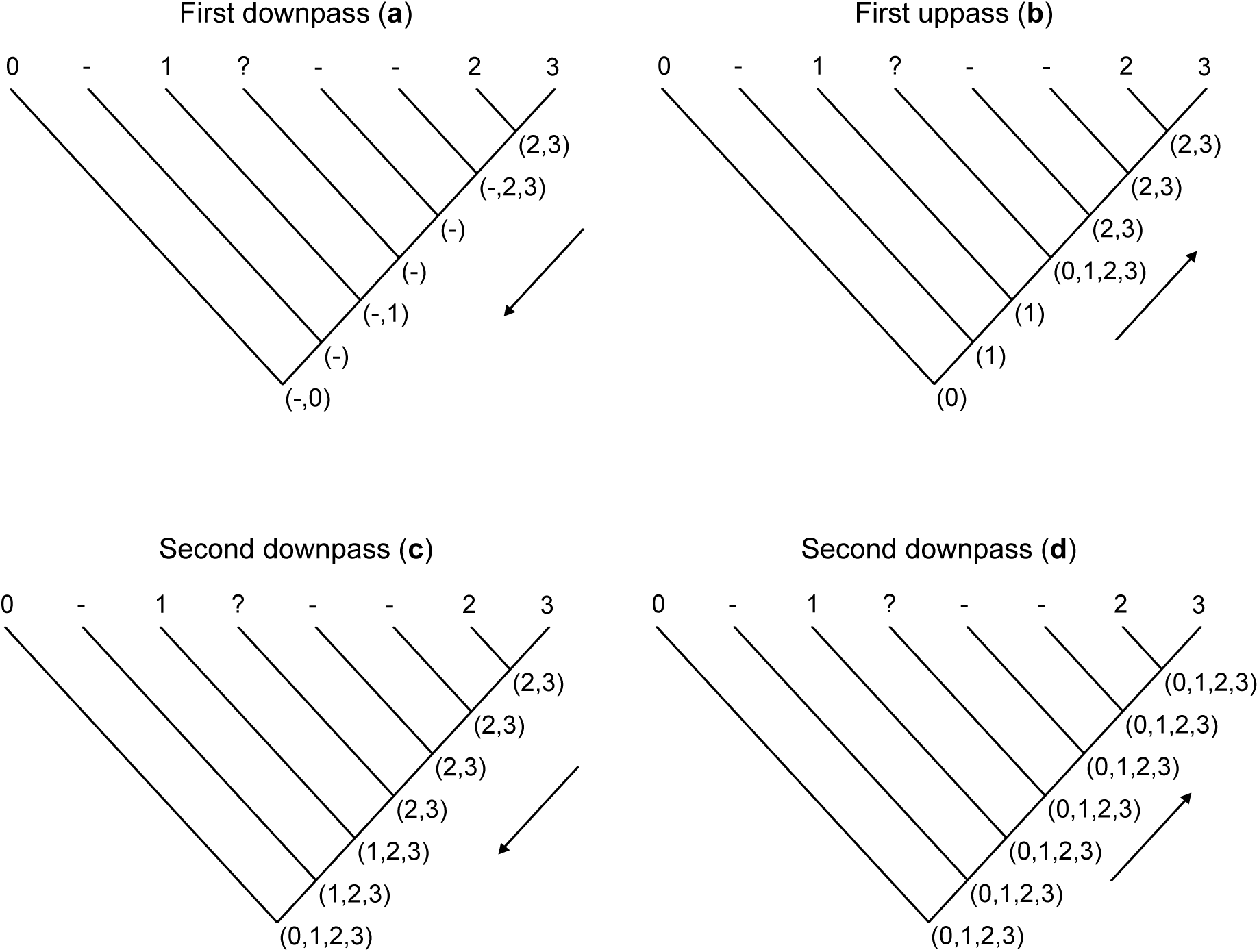
Illustration of the use of four passes for correctly estimating the ancestral states with inapplicable data in this phylogeny. After the second pass, the ancestral state sets are incorrect at six of the seven nodes. **else** *set* the node’s state to be the token in common between both descendants.

### First postorder traversal (downpass) – Figs 2a, 3a

Traverse the internal nodes of the tree in postorder. At each node:

1. **If** there is any token in common between both descendants, *go* to 2; **else** *go* to 3.
2. **If** the token in common is only the inapplicable token, and both descendants have an applicable token, *set* the node’s state to be the union of the descendants’ states;
3. **If** both descendants have an applicable token, *set* the node’s state to be the union of both descendants’ states without the inapplicable token; **else** *set* the node’s state to be the union of its descendants’ states. **Then** *go* to 4.
4. Visit the next node in postorder. Once all nodes have been visited, *conduct the first uppass*.

### First preorder traversal (uppass) – Figs 2b, 3b

Traverse the tree in preorder. At each node:

1. **If** the node has the inapplicable token, *go* to 2; **else** leave the node’s state unchanged and *go* to 8.
2. **If** the node also has an applicable token, *go* to 3; **else** *go* to 4.
3. **If** the node’s ancestor has the inapplicable token, *set* the node’s state to be the inapplicable token only and *go* to 8; **else** *remove* the inapplicable token from the current node’s state. **Then** *go* to 8.
4. **If** the node’s ancestor has the inapplicable token, *set* the node’s state to be the inapplicable token only and *go* to 8; **else** *go* to 5.
5. **If** any of the descendants have an applicable token, *set* the node’s state to be the union of the applicable states of its descendants; **else** *set* the node’s state to be the inapplicable token only. **Then** *go* to 8.
6. **If** the unvisited tip includes both inapplicable and applicable tokens, *go* to 7; **else**
7. **If** the current node has only the inapplicable token, *set* the tip’s state to the inapplicable token only; **else** *remove* the inapplicable token from the tip’s state. **Then** *go* to 8.
8. **If** one of the node’s descendants is an unvisited tip, *go* to 6; **else** visit the next node in preorder. Once all nodes and tips have been visited, *initialise the tracker*.

### Initialise tracker – Figs. 2b, 3c

Visit each tip in turn. At each tip:

1. **If** the tip’s state contains the inapplicable token, *set* its tracker to “off” and *go* to 4;
2. **else** *go* to 2.
3. **If** the tip’s state does not contain the inapplicable token, *set* its tracker to “on” and *go* to 4; **else** *go* to 3.
4. **If** the ancestor’s state contains an inapplicable token, *set* the tip’s tracker to “off”;
5. **else** *set* the tip’s tracker to “on”. **Then** *go* to 4. D. Visit the next tip. Once all tips have been visited, *conduct the second downpass*.

### Second postorder traversal (downpass) – Figs. 2b, 3c

Traverse the tree in postorder. At each node:

1. **If** the tracker of either descendant is “on”, *set* this node’s tracker to “on”; **else** *set it* to “off”. **Then**, *go* to 2
2. **If** the node had an applicable token in the first uppass, *go* to 3; **else** leave the node’s state unchanged and *go* to 8.
3. **If** there is any token in common between both descendants, *go* to 4; **else** *go* to 5.
4. **If** the tokens in common are applicable, *set* the node’s state to be the tokens held in common, without the inapplicable token; **else** *set* the node’s state to be the inapplicable token. **Then** *go* to 8.
5. *Set* the node’s state to be the union of the states of both descendants (if present) without the inapplicable token, and *go* to 6.
6. **If** both descendants have an applicable token, ***increment*** the tree score (step increment) and *go* to 8; **else** *go* to 7.
7. **If** both of the node’s descendants’ trackers are “on”, ***increment*** the tree score (applicable region increment) and *go* to 8; **else** just *go* to 8.
8. Visit the next node in postorder. Once all nodes have been visited, *conduct the second uppass*.

### Second preorder traversal (uppass) – Figs. 2b, 3d

Traverse the tree in preorder. At each node:

1. **If** the node has any applicable token, *go* to 2; **else** *go* to 9.
2. **If** the node’s ancestor has any applicable token, *go* to 3; **else** *go* to 10.
3. **If** the node’s state is the same as its ancestor’s, *go* to 10; **else** *go* to 4.
4. **If** there is any token in common between the node’s descendants, *go* to 5; **else** *go* to 6.
5. *Add* to the current node’s state any token in common between its ancestor and its descendants and *go* to 10.
6. **If** the states of the node’s descendants both contain the inapplicable token, *go* to 7; **else** *go* to 8.
7. **If** there is any token in common between either of the node’s descendants and its ancestor, *set* the node’s state to be its ancestor’s state; **else** *set* the current node’s state to be all applicable tokens that are common to both its descendants and ancestor. **Then** *go* to 10.
8. *Add* to the node’s state the tokens of its ancestor. **Then** *go* to 10.
9. **If** both of the node’s descendants’ trackers are “on”, ***increment*** the tree score (applicable region increment) and *go* to 8; **else** *go* to 10.
10. Visit the next node in preorder. Once all nodes have been visited, *calculate the tree score*.

### Calculate tree score

The contribution of the given character to the total score of the tree is given by:

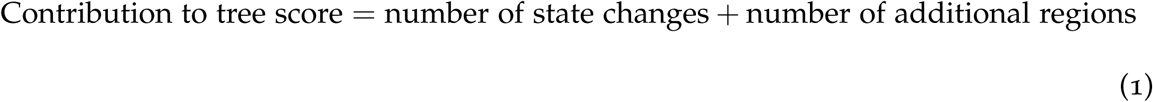

State changes are recorded in the second downpass (at point 6); the number of applicable regions is calculated in both the second downpass (at point 7) and the second uppass (point 9).

## Character coding with inapplicable-aware reconstruction algorithms

### Two categories of ontologically dependent character

An upshot of recognizing maximized homology and minimized homoplasy as the objective of maximum parsimony is that not all cases of ontological dependency of characters (*sensu* Vogt, 2017) require reductive coding. Two coding strategies may be applied, depending on the *information* implied by the states of an ontologically dependent character. That is, we may distinguish between subcharacters that are *transformational* (as in the case of tail colour) and *neomorphic* (following Sereno, 2007).

Transformational character statements describe a variable property of a principal character, with no biological reason to anticipate any particular ancestral state. The case of tail colour, as discussed above, is transformational; with reductive coding, it can be correctly handled by our algorithm. If a red tail appears twice on a particular tree topology, then the fact that it is red in both instances represents an instance of homoplasy: an independent innovation of the colour red. Using the inapplicable token to denote tail colour in non-tail-bearing taxa (Table 1) causes our algorithm to recognize the second innovation of a red colouration as an instance of homoplasy that should contribute to the tree’s length.

Neomorphic character statements are presence/absence characters that depend on the presence of the principal character. An example would be the presence of eyespots on a tail. Such characters may be scored as binary characters without the use of the inapplicable token, as long as there is still a separate character for presence/absence of tail. Given the presence of a tail, a researcher might conclude that the absence of eyespots, or equivalent features, is unsurprising. Two separate instances of a tail without eyespots would then be said to exhibit a homoplasy with respect to tail presence, but not with respect to the absence of eyepsots. Unlike the case of tail colour (a tail must have *some* colour when it appears), the presence of an eyespot is not necessarily expected. The absence is an uninformative value, and therefore would be more difficult to describe as homoplasious. If, on the other hand, eyespots are present on the two occasions that a tail appears, then the second occurrence of eyespots does represent an instance of homoplasy. Likewise, a secondary *loss* of eyespots elsewhere on the tree would represent an instance of convergence and should therefore contribute to tree length. For this reason, simple binary presence/absence coding may be employed, where an absence value would cover both observed absence and absence due to the absence of the principal character (Table 1). Even when applying our algorithm, inapplicable tokens should not be employed in such instances, as they would incorrectly penalize trees in which a tail (lacking eyespots) originated multiple times. If additive characters are decomposed into a series of neomorphic characters, then the original and decomposed characters are mathematically equivalent under this coding approach.

**Table 1:** Coding inapplicable data in ontologically dependent characters. “Tail” is a principal character with two ontologically dependent characters, “Tail colour”, a transformational character that should be coded as “-” when a tail is absent, and “Tail eyespot”, a neomorphic character that should be coded as “0” when a tail is absent.

|  |  |  |  |  |  |  |  |  |  |
| --- | --- | --- | --- | --- | --- | --- | --- | --- | --- |
| Tail | 0 | 0 | 0 | 1 | 1 | 1 | 1 | 1 | (0, absent; 1, present) |
| Tail colour | - | - | - | 0 | 0 | 1 | 2 | ? | (0, red; 1, blue; 2, green; -, inapplicable) |
| Tail eyespot | 0 | 0 | 0 | 1 | 0 | 1 | 0 | ? | (0, absent; 1, present) |

### “Parsimony-uninformative” characters inform parsimony

A consequence of our approach is that the distribution of inapplicable tokens conveys grouping information. A topology that implies that red tail colouration evolved once has a shorter length than one on which red tail colouration evolved twice, even if no other colour of tail is observed. This seemingly counterintuitive result arises because the algorithm prefers trees that attribute similarities to common ancestry rather than to chance. The effect of including ontologically dependent characters that would not be parsimony-informative under the standard Fitch algorithm is to up-weight the corresponding principal character. Care must be taken, therefore, that each ontologically dependent character truly reflects a biologically significant similarity, for a principal character might be misleadingly upweighted if trivial subordinate properties (e.g. “number of DNA bases in tail”) are included in a matrix.

### Missing but applicable character states

With an implementation of the Fitch algorithm that does not consider the inapplicable token as equivalent to missing data, extra care should be employed when coding missing data. Consider an example dataset that includes fossil and extant species.

Following the character “Tail colour” described above (Table 1), one could code a fossil taxon where the tail is entirely missing due to incomplete preservation. In this case, the “Tail colour” should be coded as “?” (it is uncertain whether the tail was red, blue or green or whether the tail was present at all). If we now consider another fossil taxon were the tail is clearly preserved but the colour is not observable, the character state ambiguity could be coded in one of three ways:

- If, as is the usual case, there is no *a priori* information indicating whether or not a tail is homologous with those of other taxa, tail colour should be coded as uncertain (“?”, treated by the algorithm as (-012)).
- If the tail of the fossil taxon is known to be homologous with the tails of other taxa, then an optimal character reconstruction will assert that its colour is one of the colours that has been observed in other taxa (the ambiguity should be coded as “red, blue or green” (012)).
- If the tail of the fossil taxon is known *not* to be homologous with the tails of other taxa, then an optimal character reconstruction will assert that its colour is *different* from any colour that has been observed, because a second innovation of a colouration observed elsewhere on the tree would represent an instance of homoplasy. In this case, tail colour should be coded as inapplicable (-), the character’s definition being effectively “colour of tail of homologous type”.

## Comparing approaches to phylogenetic reconstruction

In order to evaluate how this approach impacts phylogenetic results, we analyzed 30 discrete morphological matrices under three approaches: (i), reductively coded datasets treated under traditional Fitch parsimony, with inapplicable treated as missing (here termed the “missing” approach); (ii), the “extra state” approach, using compound coding with inapplicability as a separate state; and (iii), the “inapplicable” approach, which applies our new algorithm.

Before beginning our analysis, every inapplicable token in each neomorphic character was replaced with the token corresponding to the presumed non-derived condition (typically “absent”). Each matrix was then subjected to phylogenetic tree search: the “missing” and “extra state” approaches used TNT, employing the parsimony ratchet, sectorial search and tree drifting algorithms (Goloboff, 1999; Nixon, 1999; Goloboff and Catalano, 2016); the inapplicable approach used R (R Core Team, 2017) for tree search, using the parsimony ratchet as deployed in our new package inapplicable (see Implementation section below). Although this latter search approach is inefficient, it nevertheless converges on an optimal tree length within minutes (<50 tips) to hours (<80 tips). Whilst it is difficult to guarantee that every optimal tree will be identified, we ensured a wide sampling of tree space by conducting 100 independent tree searches in TNT, and by sampling shortest trees in R until the shortest length had been found by 250 ratchet iterations.

In order to establish whether the three methods recovered different sets of optimal trees, the number of trees that occurred in the optimal sets of one, two, or all three approaches was tallied. In addition, a strict consensus tree was calculated for all trees in each optimal set, the number of bipartitions present in each set serving as a proxy for the disparity of trees that is optimal under each approach. Finally, each set of optimal trees was plotted in a two-dimensional space (Hillis et al., 2005) by decomposing a matrix of pairwise quartet distances (Estabrook et al., 1985), calculated using the tqDist R library (Sand et al., 2014), into two dimensions by minimising the Kruskal-1 stress function (Borg and Groenen, 2005), following Hillis et al. (2005).

## Results

In most cases, the three different methods identified different sets of optimal trees. Indeed, only in one of the thirty examined datasets were the optimal trees recovered by each method also optimal under the other two (Fig. 4a). In ten datasets (Fig. 4b), a subset of trees are optimal under all methods, but other trees are optimal under one method and a few steps longer under another. In nine datasets (Fig. 4c), the forests of trees that are optimal under two methods (here, “missing” and “extra state”) partially overlap, but in one method (here, “inapplicable”), no optimal trees were found that are also optimal under either other method. In the final ten datasets (Fig. 4d), each method generates a distinct set of optimal trees. Summing across all datasets, only 4% of trees that were optimal under one method were also optimal under the other two (Fig. 5a).

**Figure 4:**
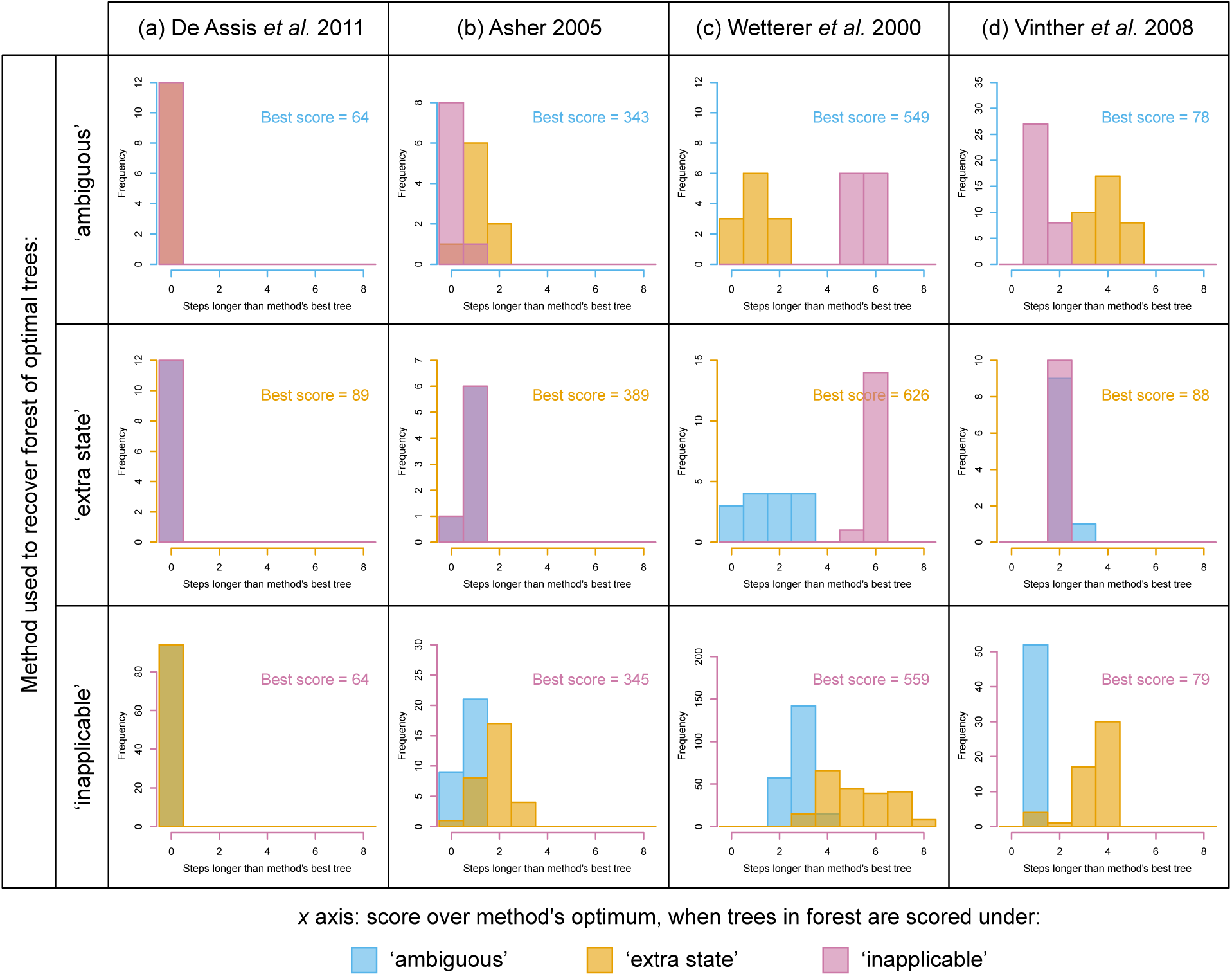
Different methods recover different optimal tree sets. Each histogram details the distribution of tree scores when a each of the optimal trees recovered under method P is scored using method Q. Scores are presented relative to the lowest score recovered by method Q for each dataset. Histograms for all examined datasets are presented in the supplementary information.

**Figure 5:**
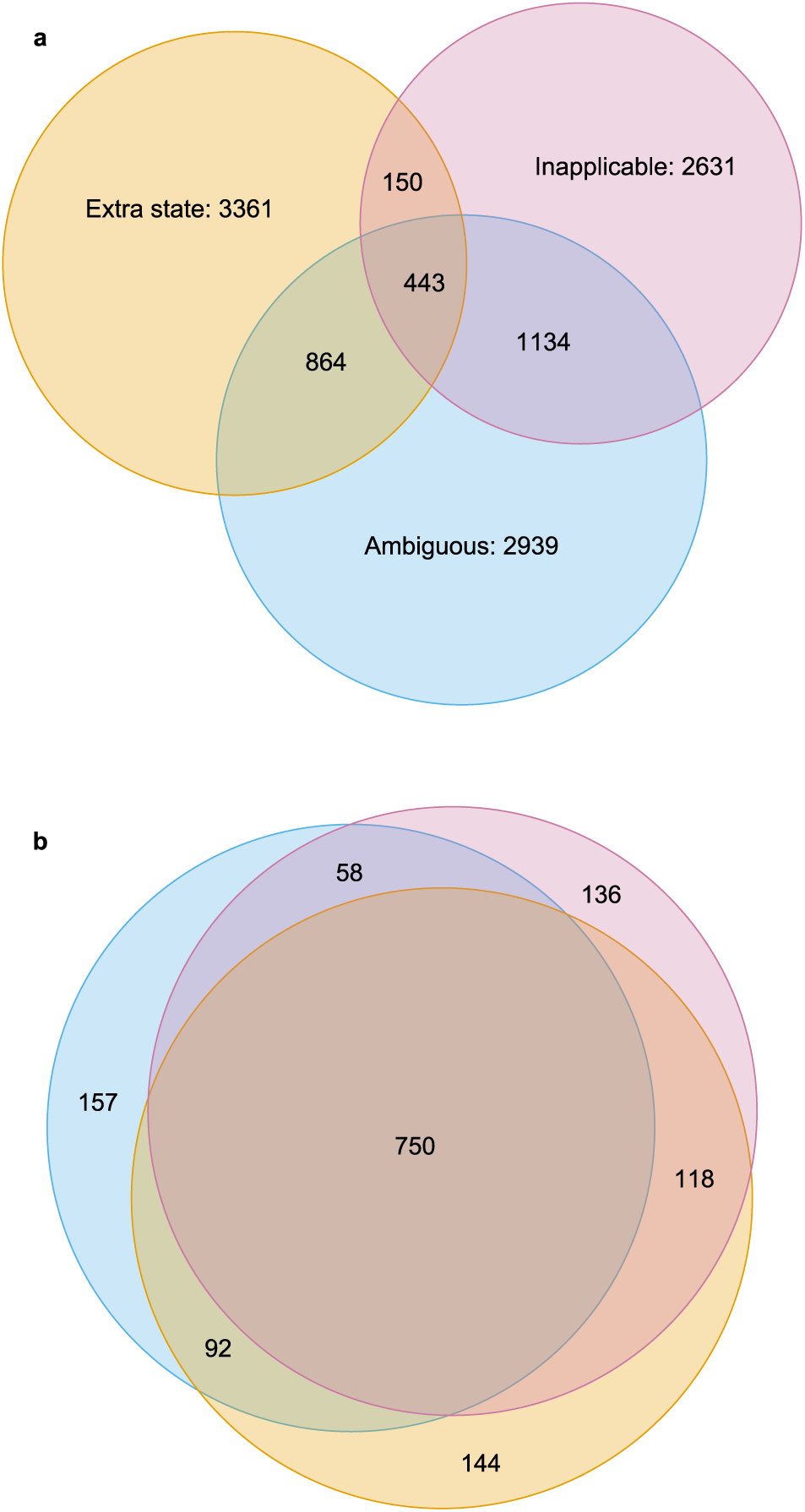
Venn diagrams depicting (a), proportions of optimal trees that are optimal under one, two or three methods; (b), proportion of nodes present in every optimal tree recovered under one, two or three methods. Results are summed across all datasets; figures for individual datasets are available in the supplementary information.

How topologically different were the trees that each method described as optimal? One qualitative way to explore the difference between multiple forests of trees is to generate a two-dimensional treespace from the distances between pairs of trees. This approach demonstrates that it is difficult to predict which methods will identify the most similar sets of optimal trees, and that the regions of treespace identified as optimal by the different methods may be very different or very similar (Fig. 6).

**Figure 6:**
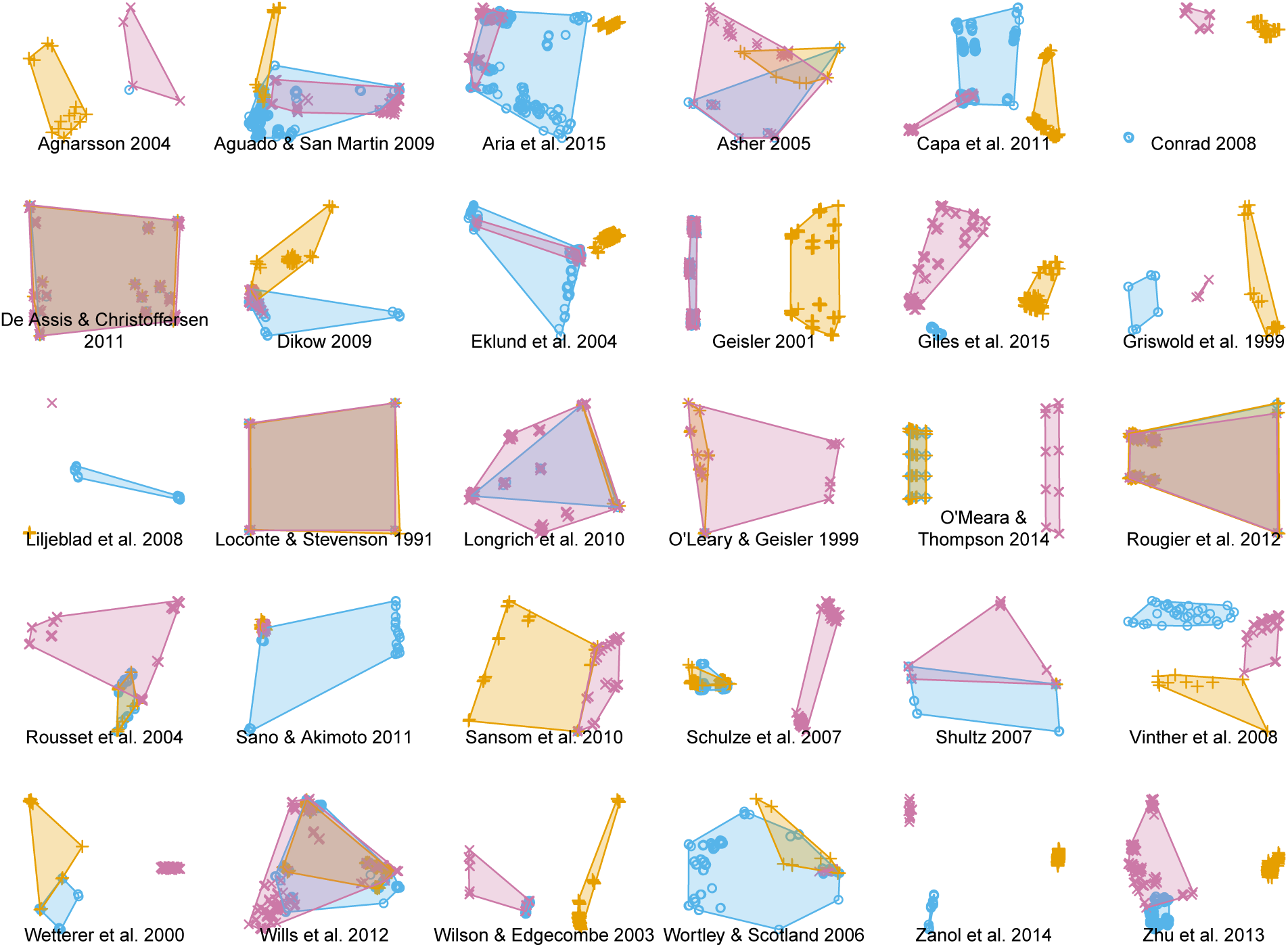
Distribution of optimal trees in MDS treespace for each dataset. Shaded regions correspond to convex hulls surrounding all optimal trees recovered using a given approach. No method is consistently more precise or more similar to any other method.

An alternative way to explore how much trees in the three optimal sets have in common is to count the number of nodes held in common between trees within a set – or, in other words, the number of nodes present on the strict consensus of all trees in that set. On this approach, averaged across all datasets, 76% of the nodes that are present in every tree that is optimal under the “inapplicable” approach are also present in every tree that is optimal under the “missing” approach, and 82% are present in every tree that is optimal under the extra-state approach; only 70% are present in all trees recovered by all methods (Fig. 5b).

Even though, in any one dataset, the number of trees identified as optimal can vary considerably between the three methods, we were unable to identify any systematic trend in the disparity of optimal trees. Neither the number of distinct trees in the optimal tree set, the resolution of the strict consensus tree, nor the area of treespace occupied by the trees showed any systematic variation.

### Implications

The accuracy of a method measures whether the method will reconstruct the “true” tree from a given dataset. As the “true” evolutionary tree is unknown, attempts to measure the accuracy of phylogenetic methods rely on data simulated from a predetermined tree topology. In the absence of a robust and testable model that can realistically simulate inapplicable morphological data, it is not possible to objectively compare the accuracy of different approaches. This said, the fact that the three approaches each identify different trees as optimal indicates that the methods differ in their accuracy. Because the “inapplicable” approach does not incorporate the errors that accompany the other approaches, we suggest that its results are the most likely to be phylogenetically accurate. This is not the same as claiming that this method improves the statistical consistency of parsimony, as it makes the same basic probabilstic assumptions. Rather, it eliminates a deficiency of parsimony methods as they are applied in practice.

The precision of an approach is more readily quantified; it represents a measure of the number or variety of trees that have an equal length under a particular counting regime. Our inapplicable method proves to be more precise than the other approaches as often as it is less precise, meaning that any improvement or loss of accuracy associated with the method comes with no effect on the precision or resolution of results.

## Conclusion

We have presented a single-character modified Fitch algorithm for ancestral state reconstructions that is aware of a special “inapplicable” token. This algorithm correctly reconstructs ancestral states by acknowledging that applicable state distributions rely on the prior resolution of applicable/inapplicable dichotomies. Because applicable state assignments depend on the resolution of the outcome of applicable/inapplicable relations, up to four passes may be required to correctly calculate tree length. Furthermore, missing data need to be updated at the tips – initially as either applicable or inapplicable – in order to complete ancestral state sequences. Our tree scoring procedure follows De Laet (2015) in penalizing increasing amounts of homoplasy without redundant penalties. Up to three traversals are necessary in order to count the number of transformations on a tree, which can be achieved during the second downpass. However, a final estimate of the number of regions on the tree is counted on the fourth traversal (final uppass). The method, unsurprisingly, takes additional time, though this is expected to be mostly in proportion to the number of characters having inapplicable tokens. Nevertheless, some economies are possible, because only characters with three or more inapplicable tokens need to be treated with this algorithm. The method provides a means of evaluating existing datasets with minimal modification, and without a need to specify explicit relationships between characters (because presence/absence information is already implicit in the applicable/inapplicable distinction). Preliminary results show that analyses with large amounts of inapplicable data are likely to be considerably affected by inapplicable data. In some cases, the set of trees that are optimal under our new algorithm does not overlap with the optimal sets obtained by existing approaches, suggesting that our method allows a gain in accuracy with no corresponding loss of precision.

### Implementations

The algorithm described throughout this paper is implemented at different levels in different projects. The main C implementation of the algorithm and associated tools is available at http://www.morphyproject.org/. An R implementation based on the former is available in the inapplicable package at https://github.com/ms609/inapplicable. Finally, a shiny (R) visualisation of the algorithm is available via the Inapp package at https://github.com/TGuillerme/Inapp. Permanent archives of the above implementations are available on FigShare, http://dx.doi.org/10.6084/m9.figshare.c.3911821

## Acknowledgements

TNT is made available with the sponsorship of the Willi Hennig Society.

